# Lung macrophages utilize unique cathepsin K-dependent phagosomal machinery to degrade intracellular collagen

**DOI:** 10.1101/2022.05.16.492085

**Authors:** Ivo Fabrik, Orsolya Bilkei-Gorzo, Daniela Fabrikova, Maria Öberg, Johannes Fuchs, Carina Sihlbom, Melker Göransson, Anetta Härtlova

## Abstract

Resident tissue macrophages (RTMs) are organ-specialized phagocytes responsible for the maintenance and protection of tissue homeostasis. It is well established that tissue diversity is reflected by the heterogeneity of RTMs origin and phenotype. However, much less is known about tissue-specific phagocytic and proteolytic macrophage functions. Here, using quantitative proteomics approach, we identify cathepsins as key determinants of phagosome maturation in primary peritoneal, lung and brain resident macrophages. The data further uncover cathepsin K (CtsK) as a molecular marker for lung phagosomes required for intracellular protein and collagen degradation. Pharmacological blockade of CtsK activity diminished phagosomal proteolysis and collagenolysis in lung resident macrophages. Furthermore, pro-fibrotic TGF-β negatively regulated CtsK-mediated phagosomal collagen degradation independently from classical endocytic proteolytic pathways. In humans, phagosomal CtsK activity was reduced in COPD lung macrophages and non-COPD lung macrophages exposed to cigarette smoke extract. Taken together, this study provides a comprehensive map of how peritoneal, lung and brain tissue environment shapes phagosomal composition, revealing CtsK as a key molecular determinant of lung phagosomes contributing to phagocytic collagen clearance in lungs.

## INTRODUCTION

Resident tissue macrophages (RTMs) are present in every organ of our body and play an essential role in host protection against infection (Bleriot, Chakarov et al., 2020). As professional phagocytes, they are specialized in the recognition, engulfment and digestion of pathogens but also apoptotic cell debris (Murray & Wynn, 2011). Thus, phagocytic process is essential not only for microbial elimination, but also for the maintenance of tissue homeostasis and remodelling (Arandjelovic & Ravichandran, 2015, Lemke, 2013, Rothlin, Ghosh et al., 2007).

Phagocytosis is a highly conserved process characterised by recognition and ingestion of particles larger than 0.5 *μ*m into a plasma membrane derived vesicle, known as phagosome (Pauwels, Trost et al., 2017). After internalization, phagosomes undergo a series of fusion steps with endosomal compartment and ultimately with lysosome to form a highly acidic phagolysosome. During this last step, the phagolysosome acquires important components for the degradation of particle, including proteases, nucleases and lipases required for digestion of engulfed material (Desjardins, Huber et al., 1994, Fairn & Grinstein, 2012). This process needs to be tightly regulated for phagocytes to carry out their function in immunity and homeostasis. If uncontrolled, it can lead to the development of persistent infection or autoimmunity and inflammatory disorders (Colegio, Chu et al., 2014, Johnson & Newby, 2009, Nagata, Hanayama et al., 2010).

It is now well established that the majority of RTMs are derived from embryonic yolk sac and fetal liver precursors independently of the contribution of bone marrow precursors (Bleriot et al., 2020, Ginhoux & Guilliams, 2016). As such, specific macrophage responses to a phagocytic prey are not only defined by utilization of phagocytic receptor and activation of distinct signalling pathways but also by microenvironment-dependent signals. However, the mechanisms by which tissue milieu impacts macrophage phagocytic capacity and downstream phagosomal processing are poorly understood. We hypothesized that different tissue environment will modulate the phagosomal system of distinct RTMs and will contribute to their organ-specific function.

Here, by employing high-resolution mass spectrometry-based proteomics, we describe differences in the phagosomal environment of primary macrophages from distinct tissues. Our data reveal an essential role of cathepsin proteases as drivers of RTMs-specific phagosomal proteolytic activity. Specifically, we identify cathepsin K as a phagosomal marker of mouse and human lung macrophages with a crucial function in phagosomal proteolysis and collagen degradation. Importantly, cathepsin K-mediated phagosomal collagenolysis in lung RTMs is negatively regulated by fibrosis-related stimuli and reduced in macrophages from COPD patients. Altogether, these data uncover that phagosomal distribution of cathepsins within distinct RTMs reflects their local substrates and determines their tissue-specific functions.

## RESULTS AND DISCUSSION

### RTMs phagosomes mature at distinct rate

We first investigated whether tissue residency impacts macrophage phagocytic activity. RTMs isolated for phagocytic profiling included brain, lung, and peritoneal macrophages. These three different RTMs represent cells of distinct macrophage origin with a specialized function essential for their tissue environment. While brain macrophages are mainly homeostatic cells providing the main cellular defense of the central nervous system, lung macrophages maintain lung physiology by clearance of pulmonary surfactant and eliminating microorganisms or allergens (Guilliams, De Kleer et al., 2013, Li & Barres, 2018). Peritoneal macrophages are mainly important for protection against pathogens (Cassado Ados, D’Imperio Lima et al., 2015). Distinct macrophage populations were isolated as described in Materials and Methods and their phenotypes were confirmed by flow cytometry (Fig. EV1). To determine phagocytic activity of different RTMs, we examined the uptake of apoptotic cells, necrotic cells, and carboxylate beads which serve as surrogates to apoptotic cells (Fig. 1A-C). Apoptosis was induced by treatment with cycloheximide while necrotic cells were induced by repeated freeze/thaw cycles. Despite distinct phagocytic pathways preferred by different RTMs (N, Quintana et al., 2017), our analysis did not reveal notable differences in dead cell or carboxylated beads uptake among RTMs. This suggests similar rates of cell debris removal by different RTMs under homeostatic conditions; probably due to functional redundancy of RTM-specific phagocytic components.

**Figure 1:**
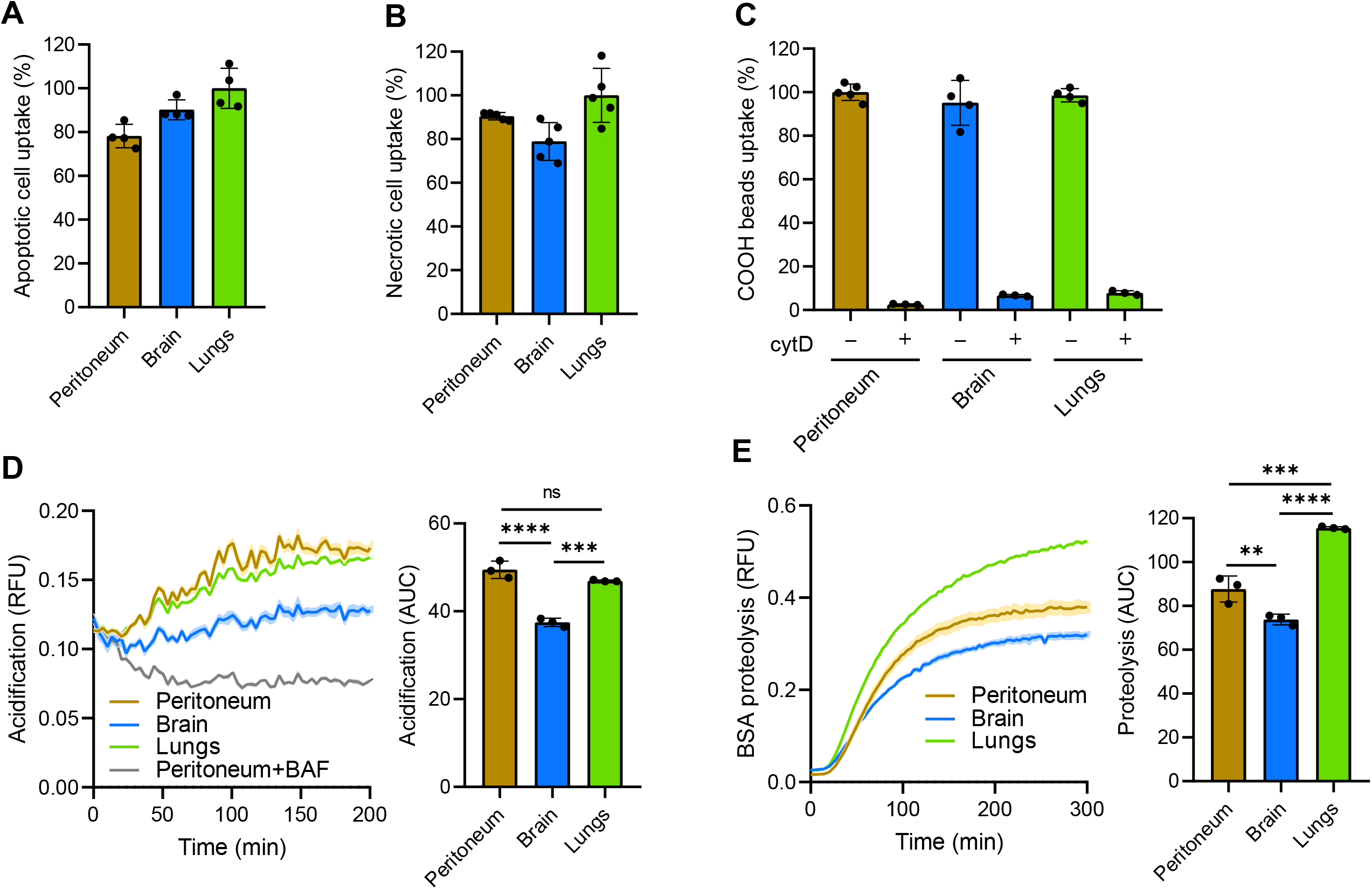
Phagosomes of RTMs mature at different rates. A-C: Uptake of apoptotic cells (A), necrotic cells (B), and BSA-coated negatively charged silica particles (C) by RTMs isolated from peritoneum, brain, and lungs. Inhibition of phagocytosis by 10 µM cytochalasin D was done 1 h before and during the experiment. D,E: Kinetic measurement of phagosomal acidification (D) and proteolysis (E) in peritoneal, brain, and lung RTMs exposed to BSA-coated particles. Inhibition of phagosomal acidification by 100 nM bafilomycin A1 was done 1 h before and during the experiment. Data information: The statistical significance of data is denoted on graphs by asterisks where ** *p*<0.01, *** *p*<0.001, **** *p*<0.0001 or ns = not significant as determined by ANOVA with post-hoc Tukey test. Data in bar graphs (A-C) are shown as means of normalized fluorescence ± standard deviation (SD). Data showing time-dependent fluorescence (D,E) are normalized to beads uptake (relative fluorescence units, RFU) and expressed as means of RFU ± standard error of the mean (SEM) or as area under curve (AUC) ± SD. Data are representative of two (D) or three (A-C, E) replicates.

Next we decided to determine whether phagosome maturation varies in distinct RTMs by analysing phagosomal pH and proteolysis in real time (Podinovskaia, VanderVen et al., 2013, Yates, Hermetter et al., 2007). While peritoneal and lung macrophages exhibited comparable rates and extents of phagosomal acidification, brain macrophages profoundly showed a smaller proportion of acidified phagosomes indicating less degradative capacity of brain phagosomes compared to lung and peritoneum (Fig. 1D). Notably, the proteolytic efficiency of lung phagosomes was significantly enhanced compared to peritoneal macrophages in spite of similar rate of phagosomal acidification. These data could be explained by a faster fusion of lung phagosomes with lysosomes and/or by a higher concentration of proteases in lysosomes of lung macrophages compared to peritoneal macrophages (Fig. 1E). Taken together, these data suggest that phagosome maturation contributes to functional heterogeneity of RTMs.

### RTMs phagosome proteome

To better understand organ-specific phagosome maturation of macrophages in more detail, we further investigated which of the phagosome-associated molecules regulate the proteolytic activity of phagosomes across different tissues. Phagosomes of peritoneal, lung and brain resident macrophages were isolated using magnetic beads and phagosomal proteomes were analysed by label free quantitative tandem mass spectrometry (Fig. 2A; Table EV1). Phagosome purity was confirmed by immunoblot showing enrichment of a prototypical phagosome marker, Rab7b, and phagosome-lysosomal marker, LAMP1, within phagosome fraction while cytosolic protein, tubulin, was absent (Fig. 2B). In line with the immunoblot data, gene ontology enrichment analysis of proteomic data confirmed enrichment of proteins involved in phagocytosis and endocytosis in the phagosomal fraction compared to low abundant cytosolic and nuclear contaminants (Fig. 2C).

**Figure 2:**
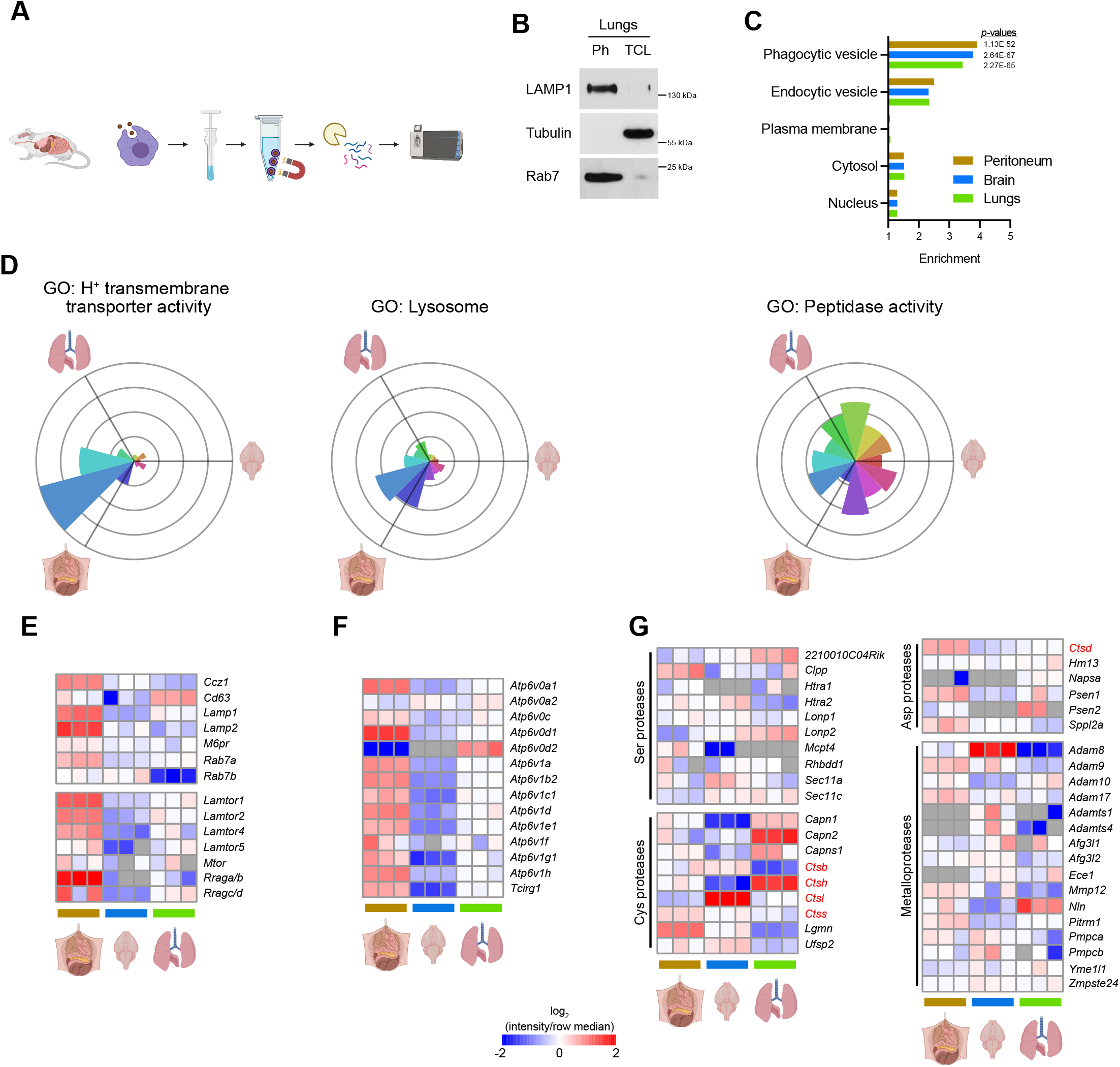
Proteins involved in phagosome-lysosome fusion are not the key determinants of phagosomal proteolytic activity. A: Workflow for isolation of phagosomes from peritoneal, brain, and lung RTMs for proteomics. B: Purity of isolated phagosomes as determined by Western blot. Presence of phagosomal markers LAMP1 and Rab7 was probed in phagosome samples (Ph) and corresponding total cell lysates (TCL). Samples from lung RTMs are shown for demonstration. C: Enrichment of phagosome-related Gene Ontology Cellular Component (GOCC) terms in RTM phagosome dataset. D: Relative distribution of proteins connected to phagosomal maturation across phagosomes of peritoneal, brain, and lung RTMs. Radius of each sector in rose plot corresponds to a relative upregulation of associated proteins in the direction of the given tissue. E-G: Heatmaps showing relative expression of components of lysosomal markers (upper part) and proteins involved in mTORC1 signalling from phagosome (E), vacuolar ATPase (F), and endopeptidases (G) in phagosomes of peritoneal, brain, and lung RTMs. Colour gradient correlates to log-ratio of intensity divided by row median, grey cells indicate missing values. Data information: Western blots (B) are representative from three replicates. Proteomic data (C-G) are derived from three replicates. Enrichment of GO terms (C) was determined by Fisher exact test yielding highlighted *p*-values for selected terms. Data in rose plots (D) are averaged from at least two replicates.

Phagosome-lysosomal fusion is a step-wise process in which phagocytic cells drive the maturation of nascent phagosomes to highly degradative, mature phagolysosomes. During this transition, the lumens of maturing phagosomes undergo a transformation characterized by decreasing pH, changes in ion composition and the addition of various hydrolases. RTMs displayed a marked variety in the composition of phagosomal proteome (Fig. EV2A). While peritoneal macrophages showed significant enrichment of proteins involved in phagosomal maturation, lung macrophages displayed enrichment of proteins involved in degradation extracellular matrix and brain macrophages in interferon signalling.

Peritoneal macrophages exhibited the highest number of up-regulated proteins associated with phagosome-lysosomal fusion (Fig. 2D and EV2B) including Rab7a GTPase, LAMP1, and LAMP2 (Fig. 2E). Additionally, the levels of four members of LAMTOR complex, a key player in lysosomal trafficking, were significantly enriched in peritoneal phagosomes compared to lung and brain (Fig. 2E). Though peritoneal and lung macrophages display similar rate of acidification, peritoneal macrophages showed increased abundances of eight V1 and three catalytic vATPase subunits, which are essential for phagosomal acidification (Fig. 2F). These data suggest that the classical phagosome maturation pathway is more active in peritoneal macrophages compared to lung and brain macrophages. Interestingly, all RTMs displayed heterogeneous expression of phagosomal proteolytic enzymes independently of proteins mediating phagosome-lysosomal fusion (Fig. 2D). These included aspartic, cysteine, serine proteases, and metalloproteases such as calpains 1 and 2, legumain, ADAM8, or ADAMTS1/4 with distinct distribution in RTM phagosomes (Fig. 2G). Of these, cathepsins showed the most consistent RTM-specific expression (Fig. 2G).

### Cathepsins are key determinants of functional specialization of RTMs

Cathepsins are the most abundant proteases of lysosomal system responsible for degradation of engulfed protein material (Bird, Trapani et al., 2009, Turk, Stoka et al., 2012). They are synthesised as inactive proenzymes and can be classified based on their catalytic site into three groups – aspartic, serine, and cysteine proteases. However, their functional relevance to distinct RTMs is still not well understood. The majority of the identified phagosomal cathepsins belong to cysteine proteases while the most abundant and ubiquitously present in all three different phagosomes was the best studied aspartic protease, cathepsin D (CtsD) (Fig. 3A). Furthermore, the proteomic analysis revealed that CtsB, C, D, S and Z were distributed across all RTM phagosomes, whereas CtsA, H, K and L showed RTM-specific phagosomal expression (Fig. 3B).

**Figure 3:**
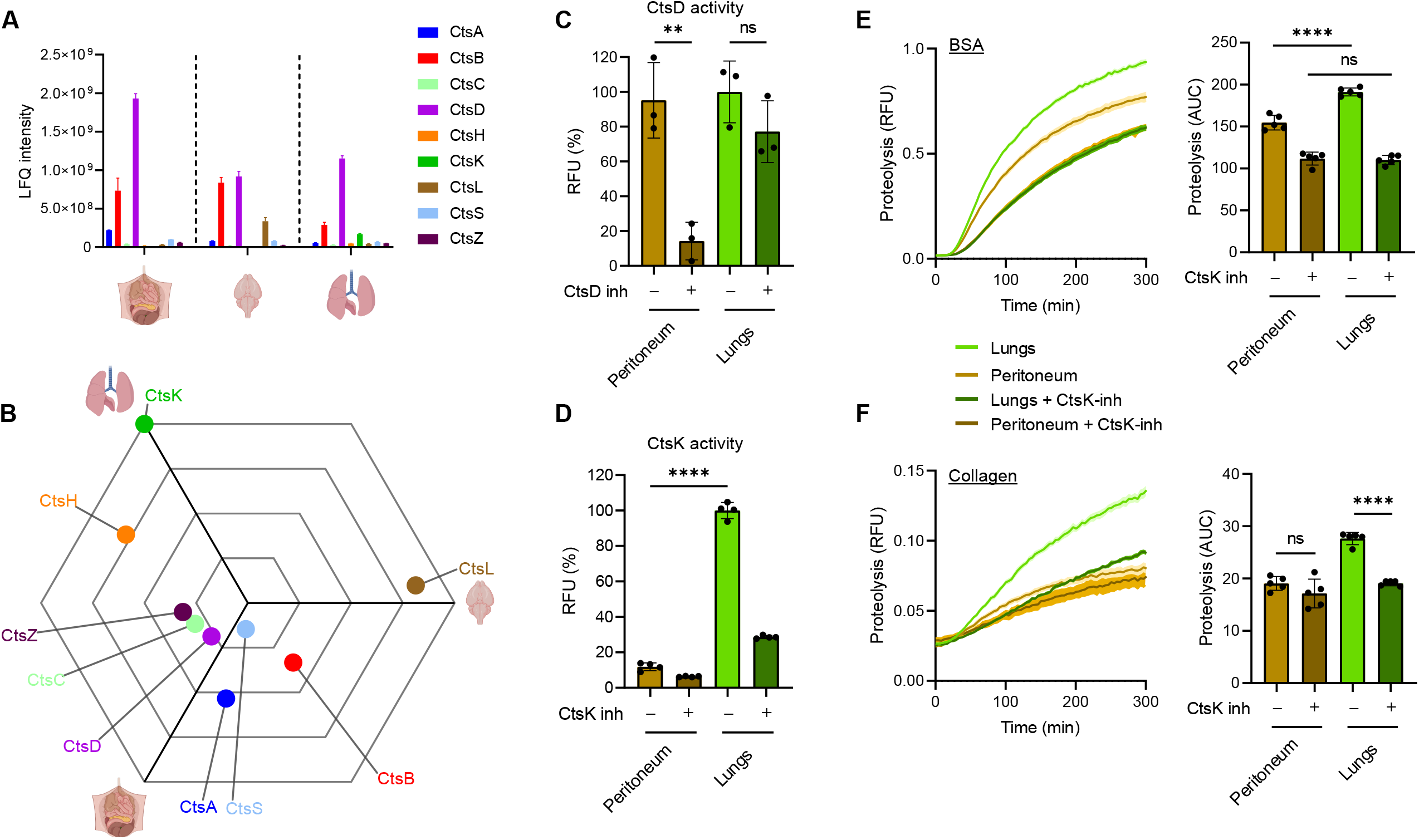
Cathepsin K enriched in phagosomes of lung RTMs is responsible for phagosomal degradation of collagen. A: Normalized LFQ intensities of individual cathepsins in RTM phagosomes as determined by MS. B: Relative distribution of cathepsins in RTM phagosomes. Distance from the centre of triwise plot corresponds to a relative upregulation of cathepsins in the direction of the given tissue. C, D: Activity of cathepsin D (C) and cathepsin K (D) in peritoneal and lung RTMs. Measured from lysates; pepstatin A (CtsD inh) and L 006235 (CtsK inh) were used at 1 µM and 10µM concentrations, respectively, and were added directly into lysates. E, F: Kinetic measurement of phagosomal BSA (E) and collagen (F) degradation in peritoneal and lung RTMs treated by CtsK inhibitor. Inhibition of CtsK by 1 µM L 006235 (CtsK inh) was done 1 h before and during the experiment. Data information: The statistical significance of data is denoted on graphs by asterisks where ** *p*<0.01, **** *p*<0.0001 or ns = not significant as determined by ANOVA with post-hoc Tukey test. Data in (A,B) are averaged from at least two replicates. Data in bar graphs (C,D) are shown as means of normalized fluorescence ± SD. Data showing time-dependent fluorescence (E,F) are normalized to uptake of coated beads and are expressed as means of RFU ± SEM. Data are representative of two (C,E) or three (D,F) replicates.

Peritoneum-enriched CtsA is a serine-type carboxypeptidase known to be involved in processing of biologically active peptides including angiotensin I or bradykinin (Timur, Akyildiz Demir et al., 2016). This correlates with high efficiency of these cells to generate bradykinin fragment and suggests that CtsA might be the effector responsible for peptide hormone signalling exerted by peritoneal macrophages (Vietinghoff & Paegelow, 2000).

Brain-enriched CtsL has been shown to be involved in microglial processing of invariant chain and remodeling of basal lamina extracellular matrix (ECM) (Gresser, Weber et al., 2001, Gu, Kanazawa et al., 2015). Interestingly, CtsL also mediates degradation of proteins involved in neurodegeneration such as tau or progranulin (Bednarski & Lynch, 1998, Lee, Stankowski et al., 2017). Our results support the crucial role of CtsL in homeostatic maintenance of neuronal tissue (Xu, Zhang et al., 2018) and we hypothesize that high phagosomal levels of CtsL in brain resident macrophages represent specific adaptation to substrates abundant in this tissue environment.

CtsH and K showed lung-specific distribution. Notably, CtsK was the only phagosomal protease which expression was solely restricted to lung phagosomes (Fig. 3A and 3B). CtsK is one of the most efficient endogenous ECM-degrading enzymes associated mostly with osteoclasts and epithelioid cells participating in remodelling of tissue (Buhling, Reisenauer et al., 2001, Dai, Wu et al., 2020), while CtsH mediates processing of alveolar Surfactant proteins B and C (Buhling, Kouadio et al., 2011, Ueno, Linder et al., 2004). Collectively, these data demonstrate that phagosomal distribution of cathepsins in distinct RTMs reflects their local substrates and determines their tissue-specific functions.

### Cathepsin K is required for phagosomal proteolytic and collagenolytic activity in lung resident macrophages

CtsK has been shown to be associated with matrix remodelling in the lungs (Buhling, Waldburg et al., 2002, Knaapi, Kiviranta et al., 2015). However, its expression by lung macrophages has so far been controversial (Buhling, Rocken et al., 2004) (van den Brule, Misson et al., 2005) (Rapa, Volante et al., 2006). To further investigate the potential specific role of CtsK in lung resident macrophages, we measured its enzymatic activity in lung and peritoneal resident macrophages and compared it to ubiquitously present CtsD. As a result, CtsD activity was similar in the cell lysate of both RTMs (Fig. 3C) while CtsK response was specific to lung resident macrophages (Fig. 3D). These data could potentially explain the enhanced phagosomal proteolytic efficiency of lung resident macrophages compared to peritoneal macrophages (Fig. 1E). To address this question, we inhibited CtsK enzymatic activity by a specific CtsK inhibitor L 006235 and measured phagosomal proteolysis in peritoneal and lung resident macrophages. This revealed that the enhanced proteolytic efficiency of lung resident macrophages was indeed dependent on CtsK (Fig. 3E). Interestingly, inhibition of CtsK activity also slightly reduced phagosomal proteolysis in peritoneal macrophages suggesting that even small amounts of CtsK (below detection by MS) might enhance proteolytic processing (Fig. 3E). Since CtsK is one of the most potent collagenases (Dai et al., 2020), we further asked whether high activity of CtsK in lung RTMs can contribute to lung tissue remodelling by mediating clearance of phagocytosed collagen. To this end, we exposed lung and peritoneal resident macrophages to collagen coated beads and measured their phagosomal collagenolytic activity. The analysis revealed that lung resident macrophages exhibited significantly higher ability to process phagocytosed collagen compared to peritoneal macrophages (Fig. 3F). Moreover, the pharmacological inhibition of CtsK enzymatic activity impaired phagosomal collagenolysis specifically in lung resident macrophages while collagen processing in peritoneal macrophages stayed intact (Fig. 3F). Altogether, CtsK contributes essentially to phagosomal proteolytic and collagenolytic activities specifically in lung resident macrophages.

### Lung resident macrophages possess CtsK-dependent phagosomal machinery tailored for ECM disposal

If CtsK plays a fundamental role in intracellular degradation of collagen in lung resident macrophages, it is expected to undergo functional changes associated with pathological states such as fibrosis. Lung fibrosis is manifested as an excessive accumulation of connective tissue driven by TGF-β (Martinez, Collard et al., 2017) (Yue, Shan et al., 2010). TGF-β stimulates production of collagen in lung fibroblasts and inhibits ECM degradation while CtsK has in general an anti-fibrotic role (Buhling et al., 2004) (Srivastava, Steinwede et al., 2008). TGF-β has been shown to downregulate CtsK expression in lung fibroblasts (van den Brule et al., 2005). Similarly to fibroblasts, TGF-β downregulated CtsK enzymatic activity in lung resident macrophages (Fig. 4A and EV3A). To further dissect TGF-β impact on CtsK function, we measured phagosomal collagen and BSA protein degradation in lung resident macrophages stimulated or not either with pro-fibrotic TGF-β or pro-inflammatory lipopolysaccharide (LPS) stimuli. The analysis revealed that pro-fibrotic TGF-β specifically reduced degradation of collagen (Fig. 4C) without affecting phagocytosis (Fig. EV3B) while pro-inflammatory stimulation decreased proteolytic activity in general (Fig. 4B and 4C). These data indicate that TGF-β specifically impacts CtsK-mediated degradation of intracellular collagen in lung resident macrophages.

**Figure 4:**
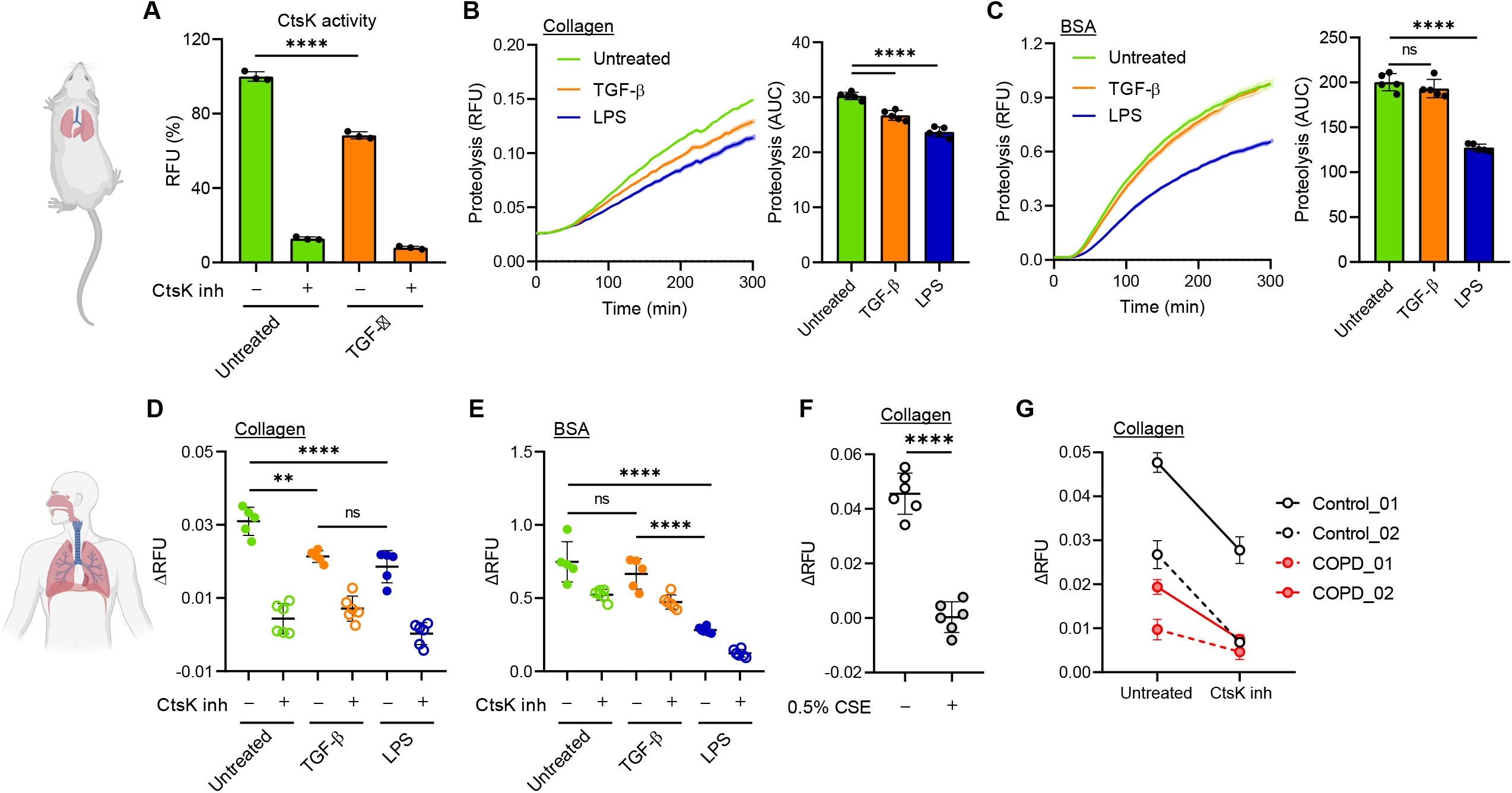
Lung RTMs possess TGF-β-regulated CtsK-dependent phagosomal machinery tailored for ECM disposal. A: Activity of cathepsin K in murine lung RTMs stimulated by TGF-β. Measured from lysates; cells were activated by murine TGF-β (5 ng/ml) 12 h before lysis. CtsK was inhibited by 10 µM L 006235 (CtsK inh) added directly into lysates. B, C: Kinetic measurement of phagosomal degradation of collagen (B) and BSA (C) in murine lung RTMs activated by TGF-β. Cells were activated by murine TGF-β (5 ng/ml) or by *E*.*coli* LPS (100 ng/ml) 12 h before the exposure to beads. D, E: Phagosomal degradation of collagen (D) and BSA (E) in human lung RTMs activated by TGF-β. Cells were activated by human TGF-β (5 ng/ml) or by *E*.*coli* LPS (100 ng/ml) 12 h before the exposure to beads. Inhibition of CtsK by 1 µM L 006235 (CtsK inh) was done 1 h before and during proteolysis. F: Inhibition of phagosomal collagen degradation by cigarette smoke extract (CSE) in human lung RTMs. Cells were treated by 0.5% CSE 12 h before exposure to beads. G: Decreased phagosomal degradation of collagen in human lung RTMs isolated from COPD patients. Inhibition of CtsK by 1 µM L 006235 (CtsK inh) was done 1 h before and during proteolysis. Data information: The statistical significance of data is denoted on graphs by asterisks where ** *p*<0.01, **** *p*<0.0001 or ns = not significant as determined by ANOVA with post-hoc Tukey test (A-E) or by Student t-test (F). Data showing time-dependent fluorescence (B,C) are normalized to uptake of collagen/BSA-coated beads and are expressed as means of RFU ± SEM or AUC ± SD. Data in (A) are shown as means of normalized fluorescence ± SD. Data in (D-G) are normalized to uptake of collagen/BSA-coated beads and are expressed as means of the fluorescence increase (ΔRFU) ± SD. Data in (A-F) are representative from three replicates.

Finally, we sought to extend the role of CtsK in phagosomal collagenolysis in human lung resident macrophages stimulated or not with TGF-β and LPS. In line with mouse data, TGF-β-stimulated lung resident macrophages exhibited significantly reduced intracellular degradation of collagen in CtsK-dependent manner while phagosomal BSA degradation was not affected (Fig. 4D and 4E). These data collectively indicate that TGF-β regulates CtsK-mediated phagosomal collagen degradation independently from classical endocytic proteolytic pathways.

### Cathepsin K-dependent collagenolytic inactivity in lung RTMs is associated with COPD pathology

The enhanced collagen clearance mediated by lung macrophages would be of critical importance during exaggerated remodelling of lung tissue, such as in chronic obstructive pulmonary disease (COPD) (Guieu & Hellon, 1980). One of the pathological hallmarks of COPD is emphysema. It is characterized by destruction of alveolar walls mediated by elastases secreted by immune cells (Abboud & Vimalanathan, 2008). This generates collagen fragments, which are pro-inflammatory and further stimulate tissue damage (Weathington, van Houwelingen et al., 2006). We hypothesized that inefficient clearance of partially digested collagen by lung resident macrophages might be another factor involved in COPD pathology. Cigarette smoke is one of the major causes associated with the development of COPD (Barnes, Burney et al., 2015). We therefore tested whether cigarette smoke extract (CSE) affects phagosomal degradation of collagen in human lung resident macrophages. CSE significantly decreased intracellular processing of collagen coated beads, even in concentrations which do not impact phagocytic activity (Fig. 4F, Fig. S3C). Next we determined collagenolytic properties of lung resident macrophages isolated from COPD patients. The analysis revealed that COPD derived lung resident macrophages exhibited reduced CtsK-dependent degradation of phagocytosed collagen compared to non-COPD control (Fig. 4G). This suggests that lung RTMs might contribute to COPD pathology by slower intracellular removal of pro-inflammatory ECM fragments.

Taken together, we show here that distribution of cathepsins in RTM phagosomes is adapted to abundant substrates of residing tissues and that lung RTMs utilize unique CtsK-dependent phagosomal pathway for clearance of phagocytosed collagen. Given that TGF-β, cigarette smoke extract, and COPD pathology all reduce CtsK-mediated collagenolysis in lung RTMs, we assume that lung tissue insults decrease the ability of lung macrophages to dispose ECM fragments and that their accumulation may exaggerate chronic lung pathologies.

## MATERIALS AND METHODS

### Isolation of murine tissue resident macrophages (RTMs)

All RTMs were isolated from 6-8 week C57BL/6 mice. All mice were maintained under specific pathogen free conditions and experiments were approved and carried out according to the guidelines set out by the Regional Animal Ethic Committee Approval #2947/20. Peritoneal RTMs were harvested by injecting phosphate buffered saline (PBS) into peritoneal cavity. Lung RTMs were obtained by homogenizing lungs using Lung Dissociation Kit (Miltenyi Biotec) followed by erythrocyte lysis. Cell suspension was then seeded into bacterial plastic dish and lung RTMs were obtained as an adherent cell fraction after overnight cultivation. Brain RTMs were isolated by homogenizing brains using Adult Brain Dissociation Kit (Miltenyi Biotec) according to manufacturer instructions. Obtained cells were enriched for CD11b^+^ cells by using CD11b MicroBeads (Miltenyi Biotec) and seeded into poly-lysine coated plastic. After isolation, all RTMs were cultivated in DMEM/F-12 with 10% fetal bovine serum (FBS), 100 U/ml penicillin, 100 µg/ml streptomycin (all Gibco), and 15% L929 supernatant for 3-5 days before using in experiments.

### Isolation of human lung macrophages

Human lung RTMs were harvested by injecting excess of PBS into human lung tissue until cell suspension started to seep out. This was repeated several times until PBS coming out from the tissue appeared completely clear. Combined cell suspension was centrifuged and erythrocytes were lysed by ACK lysis buffer (Gibco). Human RTMs were resuspended in Xvivo10 medium (Lonza) supplemented with 2 mM glutamine and penicillin/streptomycin (Gibco, 10,000 U/ml), seeded as required, and used in experiments the following day. Human lung samples were obtained from patients with or without COPD undergoing lung cancer resection surgery at the Department of Cardiothoracic Surgery at Sahlgrenska University Hospital. All human lung tissue samples were acquired in accordance with hospital and AstraZeneca ethical guidelines with written consent from all patients. Ethics committee Approval number 1026-15.

### Phagocytosis assay

RTMs were seeded into 384-well plate at a density 2.5×10^4^ cells per well 24 h before experiment. Cells were treated by 3.0 µm silica beads (Kisker Biotech) coated by AF488-conjugated BSA diluted at a ratio 1:100 by complete DMEM/F-12 medium and kept for 30 min at 37 °C. Treatment was stopped by aspirating medium followed by the addition of trypan blue to quench the fluorescence of extracellular beads. After aspirating of trypan blue, fluorescence was measured on SpectraMax i3x plate reader (Molecular Devices) using excitation/emission wavelengths of 488/525 nm. When indicated, RTMs were treated by cytochalasin D (10 µM, Sigma) for 1 h before and during the treatment.

### Phagosome functional assays

For phagosomal proteolysis assay, RTMs were seeded into 384-well plate and treated by 3.0 µm silica beads (Kisker Biotech) coated either by DQ Green BSA or DQ Collagen, type I (both Thermo). Beads were diluted at a ratio 1:100 by binding buffer (1 mM CaCl_2_, 2.7 mM KCl, 0.5 mM MgCl_2_, 5 mM dextrose, 10 mM HEPES and 5% FBS in PBS pH 7.2) and kept with cells for 5 min. Following treatment, beads were replaced by warm binding buffer and real-time fluorescence was measured at 37 °C on SpectraMax i3x plate reader using excitation/emission wavelengths of 470/525 nm. The experimental setup for phagosomal acidification assay was similar except for beads coated by BSA conjugated with pHrodo Red AM (Thermo) were used and the measured fluorescence was at 560/590 nm. All beads were labelled by AF647 and the reported fluorescence was corrected for beads loading by normalizing to AF647 signal (ex/em 640/665 nm). Quantitative data are presented either as area under curve (AUC) or as an increase of fluorescence in time (ΔRFU). When indicated, RTMs were treated by bafilomycin A1 (100 nM, Enzo) or L 006235 (1 µM, Tocris) for 1 h before and during the treatment and by murine TGF-β (5 ng/ml, R&D) or *E. coli* 055:B5 LPS (100 ng/ml, InvivoGen) 12 h before treatment.

### Uptake of apoptotic and necrotic cells

Mouse embryonic fibroblast were treated by 50 µM cycloheximide for 12 h to induce apoptosis or subjected to four freeze/thaw cycles to induce necrosis. Apoptotic and necrotic cells were then washed by ice cold PBS and labelled by 5 µM SYTO Red dye (Thermo) for 30 min at 4 °C in dark. After staining, cells were washed by ice cold PBS, resuspended in complete DMEM/F-12 medium, and added to RTMs seeded in 384-well plate at approximate ratio 10:1 (cells: RTM). After 4 h at 37 °C, RTMs were washed extensively by PBS and the fluorescence signal of SYTO Red was measured on SpectraMax i3x plate reader.

### Cathepsin activity assays

Cathepsin activity was determined by Cathepsin K and Cathepsin D Activity Assay Kits (both from Abcam) according to manufacturer instructions. Briefly, 1×10^6^ RTMs were washed by PBS, lysed on ice for 10 min using supplied lysis buffer, and centrifuged (13,000 g, 5 min, 4 °C). Supernatants were mixed with cathepsin-specific fluorescent substrate and optionally with 10 µM L 006235 or 1 µM pepstatin A (Tocris) and incubated for 1 h at 37 °C. Fluorescence was measured at excitation/emission wavelengths of 400/505 nm and 328/460 nm for cathepsin K and cathepsin D substrates, respectively. When indicated, RTMs were treated by murine TGF-β (5 ng/ml, R&D) 12 h before lysis.

### Cigarette smoke extract (CSE) preparation and cell treatment

Smoke from two 2R4F reference cigarettes (University of Kentucky) was passed through 25 ml of serum-free PBS for a total time of 10 min at a flow rate of 0.07 L/min. Obtained 100% CSE was filtered through 0.22 µm filters and used directly or snap-frozen and stored at -80 °C. For CSE treatment, cell cultivation medium was replaced by medium-diluted CSE as indicated and kept with cells for 12 h before further experiments.

### Flow cytometry

RTMs were collected at 5×10^5^ per FACS tube, washed by FACS buffer (1% BSA/5mM EDTA/PBS, pH 7.2), and incubated in blocking buffer (1:100 anti-CD16/CD32 in FACS buffer) for 15 min at 4 °C. After blocking, RTMs were washed by FACS buffer and stained by anti-CD45-BV711 (BD), F4/80-PE (BioLegend), CD11b-FITC (BD), SiglecF-PE (BD), CD11c-APC-Cy7 (BioLegend), or CD80-APC (BD) at a dilution of 1:200 by FACS buffer for 30 min at 4 °C. After washing by FACS buffer, cells were analysed by LSRFortessa X-20 (BD).

### Isolation of phagosomes

RTMs isolated from 5-10 mice were washed three times by PBS and then incubated for 30 min at 37 °C with 1.0 µm carboxylated magnetic beads (Dynabeads, Thermo Fisher Scientific) diluted at 1:20 ratio by complete DMEM/F-12 medium. Cells were then washed by ice-cold PBS, scraped, and cell pellets were washed two times by ice-cold PBS and once by hypotonic lysis buffer (HLB; 250 mM sucrose/3 mM imidazole, pH 7.4). Cells were lysed in HLB containing inhibitors of phosphatases and proteases (EDTA-free, Thermo Scientific) and 250 U/ml of Nuclease (Thermo Scientific) by 15 strokes in dounce homogenizer. Tubes with lysates were then transferred into magnetic holder and supernatants were removed. Phagosomes were washed four times by ice-cold PBS and centrifuged (13,000 g, 5 min, 4 °C) after final wash. Supernatant was discarded and pelleted phagosomes were stored at -80 °C.

### Phagosome sample preparation for proteomics

Phagosome pellets were dissolved in 25 µl of 5% SDS/50 mM triethylammonium bicarbonate (TEAB) buffer (pH 7.6) and centrifuged. Supernatants were transferred into new tubes and protein concentration were determined by BCA assay (Sigma). Approx. 6 µg of phagosomal proteins from each sample were reduced by 10 mM tris(2-carboxyethyl)phosphine (TCEP) for 30 min at room temperature (RT) and alkylated by 10 mM iodoacetamide (IAA) for 30 min in dark. The excess of IAA was quenched by addition of TCEP (final concentration 20 mM) for 15 min. Protein digestion was done using S-Trap (Protifi) according to manufacturer instructions. Briefly, samples were acidified by phosphoric acid, mixed with 6 volumes of 90% methanol (MeOH)/100 mM TEAB, and loaded onto S-Trap phase. Columns were then washed four times by 90% MeOH/100 mM TEAB and captured proteins were digested by 1 µg of trypsin (Pierce) in 50 mM TEAB at 37 °C overnight. After digestion, peptides were eluted by consecutive washes of 50 mM TEAB, 0.2% formic acid (FA), and 50% acetonitrile (ACN)/0.2% FA and the combined eluate was evaporated to approx. 20 µl. Samples were acidified by triflouroacetic acid (TFA), desalted using reversed-phase spin columns (Thermo Scientific), and dried on Speed-Vac.

### LC-MS/MS analysis

Easy nanoLC 1200 liquid chromatography system connected to Orbitrap Lumos Tribrid mass spectrometer (Thermo Fisher Scientific) were used for proteomic analyses. Peptides were first introduced onto trap column (PepMap 100 C18, 5 μm, 0.1×20 mm) and then separated on an in-house packed analytical column (Reprosil-Pur C18, 3 μm, 0.075×330 mm, Dr Maisch) using gradient (0.2% FA in water as phase A; 0.2% FA in ACN as phase B) running from 6 to 35% B in 167 min and from 35 to 100% B in 3 min, at a flow rate of 300 nl/min. Positive ion MS scans were acquired at a resolution of 120,000 within *m/z* range of 400-1600 using AGC target 5×10^5^ and maximum injection time 50 ms. MS/MS analysis was performed in data-dependent mode with a top speed cycle of 1 s for the most intense multiply charged precursor ions. MS precursors above 50,000 threshold were isolated by quadrupole using 0.7 *m/z* isolation window and then fragmented in ion trap by collision-induced dissociation (CID) at a collision energy of 35%. Dynamic exclusion was set to 60 s with 10 ppm tolerance. MS/MS spectra were acquired by ion trap using AGC target 5×10^5^ and maximum injection time 35 ms.

### Proteomic data search

Proteomic datasets were processed by MaxQuant ver. 1.6.10.43 (Cox & Mann, 2008). Data were searched against reference proteome of *Mus musculus* (UP000000589; August 7, 2020) downloaded from Uniprot. MaxQuant-implemented database was used for the identification of contaminants. Protein identification was done using these MaxQuant parameters as follows: mass tolerance for the first search 20 ppm, for the second search from recalibrated spectra 4.5 ppm; maximum of 2 missed cleavages; maximal charge per peptide z = 7; minimal length of peptide 7, maximal mass of peptide 4600 Da; carbamidomethylation (C) as fixed and acetylation (protein N-term) and oxidation (M) as variable modifications with the maximum number of variable modifications per peptide set to 5. Trypsin with no cleavage restriction was set as a protease. Mass tolerance for fragments in MS/MS was 0.5 Da, taking the 8 most abundant peaks per 100 Da for search (with enabled possibility of cofragmented peptide identification). FDR filtering on peptide spectrum match was 0.01 and only proteins with at least one identified unique peptide were considered further. Proteins were quantified using MaxLFQ function (Cox, Hein et al., 2014) with at least one peptide ratio required for pair-wise comparisons of protein abundance between samples. Proteins identified as contaminants were removed before any further interpretation of data.

### Data interpretation

Annotation of proteins by Gene Ontology (GO) was done in Perseus ver. 1.6.2.1 (Tyanova, Temu et al., 2016) and the enrichment was determined by Fisher exact test. For protein clustering, LFQ intensities of the given protein in each sample were divided by the median protein intensity and ratios were log-transformed. Values were clustered by K-means in R and the data were visualized by ‘pheatmap’ package employing hierarchical clustering of samples using Euclidean distance. Only proteins identified in all three replicates in each tissue were considered for the clustering. Triwise plot and rose plots were constructed using ‘triwise’ package (van de Laar, Saelens et al., 2016) in R by taking log-transformed protein LFQ intensities averaged from at least two replicates per tissue.

### Western blot

Phagosome samples were lysed in 5% SDS/50 mM TEAB and protein concentration was determined by BCA assay. For electrophoresis, samples were mixed with 4× Laemmli sample buffer and β-mercaptoethanol, heated to 95 °C for 5 min, and run on a NuPAGE 4–12% Bis‐Tris gel (Life Technologies). Proteins were transferred onto PVDF membrane using Mini Trans-Blot Cell system (Bio-Rad). Membranes were blocked by 5% non-fat dried milk in Tris buffered saline/0.1% Tween-20 (TBS-T) for 1 h at RT, incubated with primary antibodies in 3% BSA in TBS-T at 4 °C overnight, and with secondary antibodies in 5% milk/TBST for 1 h at RT. After incubation with HRP‐labelled secondary antibodies, proteins were detected using ECL and X‐ray films. Following antibodies were used: anti-LAMP1 (Cell Signaling), tubulin (Abcam), and Rab7 (Cell Signaling).

### Data availability

The mass spectrometry proteomics data have been deposited to the ProteomeXchange Consortium (http://proteomecentral.proteomexchange.org) via the PRIDE partner repository (Perez-Riverol, Csordas et al., 2019) with the dataset identifier PXD033010. Reviewers can access the data through the Username: and the Password: Rcpt0kMI.

## ACKNOWLEDGEMENTS

This work was funded by the Knut and Alice Wallenberg Foundation and the Wallenberg Centre for Molecular and Translational Medicine (dedicated to AH), University of Gothenburg, Sweden, Cancerfonden 19 0352 (dedicated to AH), Wilhelm och Martina Lundgrens Stiftelser 2020, 2021 (dedicated to AH). We would like to thank to the Core Proteomics facility at Sahlgrenska academy at the University of Gothenburg, Sweden.

## AUTOR CONTRIBUTIONS

AH conceived the study. IF designed and performed *in vitro* experiments, mass spectrometry protein analysis and bioinformatics analysis. CS, JF provided expertise in mass spectrometry-based proteomics. OBG performed *in vitro* experiments in human lung resident macrophages. MG provided the access to COPD patients, human lung macrophages and cigarette smoke extract. DF, MÖ performed *in vitro* experiments. AH and IF wrote the paper with contributions of all authors.

## Expanded view Figure legends

**Expanded View Figure EV1: Phenotypes of RTMs isolated from murine peritoneum, brain, and lungs**.

Flow cytometric characterization of isolated murine RTMs following cultivation for 3-5 days *ex vivo*. Representative results from two experiments.

**Expanded View Figure EV2: RTM-specific phagosomal proteins**.

A: Clustering of quantified proteins from phagosomes of peritoneal, brain, and lung RTMs. GO terms enriched in RTM-specific phagosomes are emphasized. Colour gradient correlates to log-ratio of intensity divided by row median.

B: Relative distribution of proteins annotated by GO terms connected to phagosomal maturation in phagosomes of peritoneal, brain, and lung RTMs.

Data information: Enrichment of GO terms in protein clusters (A) was determined by Fisher test. The statistical significance of data in (B) is denoted by asterisks where * *p*<0.05, ** *p*<0.01, and **** *p*<0.0001 as determined by ANOVA with post-hoc Tukey test. Data points in (B) are averaged from at least two replicates per tissue.

**Expanded view Figure EV3: Activation of lung RTMs by TGF-β or low doses of CSE does not impair their phagocytic activity**.

A: Downregulation of CD80 surface expression in TGF-β-activated murine lung RTMs measured by flow cytometry. Cells were activated by murine TGF-β (5 ng/ml) 12 h before staining.

B, C: The effect of TGF-β (B) and CSE (C) treatment on the phagocytic activity of isolated human lung RTMs. Measured by the uptake of carboxylated beads. Cells were activated by 5 ng/ml human TGF-β, 100 ng/ml LPS, or by different concentrations of CSE 12 h before exposure to beads.

Data information: The statistical significance of data in (B) is denoted by asterisks where * *p*<0.05, ** *p*<0.01, and *** *p*<0.001 as determined by ANOVA with post-hoc Tukey test. Data are representative from two replicates.

